# An inflammatory gene set–driven epigenetic clock tracks down disease progression and rejuvenation

**DOI:** 10.64898/2026.06.04.730043

**Authors:** Péter Sándor, Csaba Kerepesi, José Pedro Castro

## Abstract

Chronic, low-level inflammation, characterized by elevated pro-inflammatory programs, including epigenetic changes, in the absence of infection, is a major driver of aging and age-related diseases. On the other side of the spectrum, aging interventions work, at least in part, by decreasing inflammation. However, the molecular connection between epigenetic aging and inflammatory profiles in chronic diseases and rejuvenation has not been established yet. This study aimed to investigate the role of a newly described inflammatory signature gene set (ISig) in aging, previously associated with accelerated aging, in the progression of chronic diseases and rejuvenation. To achieve this, we developed inflammation-derived epigenetic aging clocks using ElasticNet regression models trained on CpG sites from ISig promoter regions. The newly developed inflammation aging clocks were validated on healthy samples and tested for their capacity to detect accelerated aging in diseased samples and rejuvenation during cellular reprogramming. The data demonstrate that the ISig inflammatory clocks accurately predict age, detect rejuvenation, and identify accelerated aging in disease contexts. Furthermore, we have demonstrated that it is possible to use a curated inflammatory gene-set with biological relevance to estimate biological age acceleration. We also developed a web application, the GeneClock Studio (available at https://ilab.sztaki.hu/geneclockstudio/), that allows researchers to apply the inflammatory aging clocks to their own DNA methylation datasets without requiring any programming expertise. Furthermore, the GeneClockStudio supports the training of new aging clocks based on an arbitrarily selected gene set in a similar way as in the case of the ISig inflammatory clocks.

## Introduction

Chronic, low-level inflammation, characterized by elevated pro-inflammatory proteins in the absence of infection, is a major driver of aging and age-related diseases. Conditions such as ischemic heart disease, stroke, diabetes mellitus, and chronic kidney disease (CKD) are among the leading causes of death worldwide^1,2^. Comprehensive analysis of systemic inflammation has proven effective in predicting cardiovascular disease, frailty, and overall mortality using inflammatory clocks such as iAge^3^. Furthermore, the aging state of the immune system, whether accelerated or decelerated, has a strong impact on the overall health and longevity trajectories, shown by the development of plasma proteomic aging clocks^4,5^. However, the possibility of using pre-selected inflammatory genes and connecting them to epigenetic age acceleration was not explored before. This study aimed to investigate whether a manually curated inflammatory gene-set had the potential to track epigenetic aging. For that purpose, we employed a collection of 127 inflammatory genes (inflammatory signature; ISig), strongly associated with transcriptomic aging, and estimated the biological age and the progression of chronic diseases, as well as rejuvenation events. Our goals were to: 1) validate ISig as an accurate epigenetic biomarker for the prediction of accelerated aging, chronic diseases, and rejuvenation, 2) show the connection between a custom inflammatory signature and epigenetic aging, and 3) enable researchers to test their own signature through a widely accessible, user-friendly application named GeneClock Studio. This application is intended to enable researchers to apply inflammatory or other aging clocks to rapidly generate models based on their own targeted gene sets.

This work uncovers the intrinsic connection between inflammatory genes and epigenetic aging, showing the ability to track chronic disease age acceleration, such as immunological disorders, and rejuvenation treatments such as OSKM partial reprogramming.

## Results

### ISig inflammatory aging clocks accurately predict age

We developed two inflammation-based epigenetic aging clocks by training machine learning models on CpG sites located in the promoter regions of ISig genes (Fig. 1). These sites were specifically selected due to their critical role in gene regulation, particularly in processes such as DNA replication and transcription, making them biologically relevant targets for modeling age-associated epigenetic changes. The inflammatory aging clocks were based on the ElasticNet regression model. The models were trained exclusively on healthy samples: Blood inflammatory clock 1 on the Hannum et al. dataset (GSE40279), and Blood inflammatory clock 2 on the ComputAgeBench training dataset. The Hannum et al. dataset (GSE40279) comprised 656 blood samples, with an age distribution that was right-skewed, spanning from 19 to 101 years, characterized by a higher concentration of younger individuals and a gradual decline toward older ages.

**Fig. 1.**
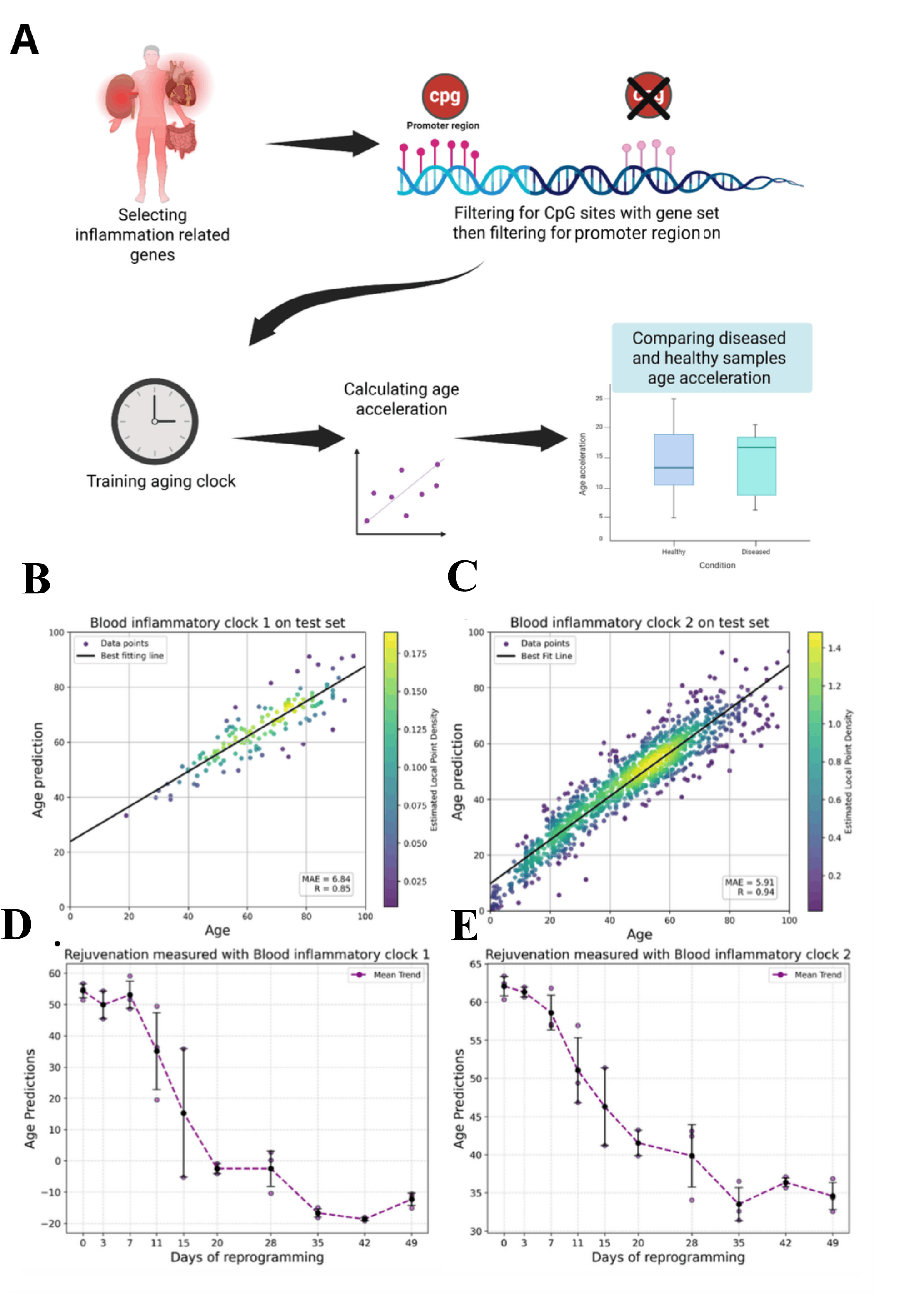
Evaluation of the ISig inflammatory clocks in aging and rejuvenation. A, Schematics of the. study workflow. An inflammation-related gene set (ISig) was first defined, and CpG sites within the promoter regions of this set were selected from the epigenetic data. Using these filtered sites, inflammation-associated epigenetic aging clocks were trained. Age acceleration was then calculated and compared between healthy and diseased samples. **B**, Predictions of the Blood Inflammatory Clock 1 on the test set, along with the best-fitting line and corresponding performance metrics. **C,** Predictions of the Blood Inflammatory Clock 2 on the test set. **D,** Predictions of the Blood Inflammatory Clock 1 on the fibroblast OSKM reprogramming dataset. **E,** Predictions of the Blood Inflammatory Clock 2 on the same dataset.

After discarding low-quality samples, the ComputAgeBench training set contained 3,180 blood samples with an approximately normal age distribution ranging from 0 to 99 years, and most samples concentrated between 30 and 60 years with a gender ratio: 50.33% female, 47.92% male, 1,75% unknown. An 80:20 training–testing split was applied for the partition of the dataset. The Blood inflammatory clock 1 performance on the test set was mean absolute error (MAE) = 6.84 years, and Pearson’s correlation coefficient (R) = 0.85 (Fig. 1B), while the Blood inflammatory clock 2 performance on the test set was MAE = 5.9 years, and R = 0.94 (Fig. 1C).

### Evaluation of the ISig inflammatory aging clock in rejuvenation

Next, we tested whether the clocks could be applied to rejuvenation strategies, shown to shut down inflammation during OSKM reprogramming^6^. In the Ohnuki et al. cell reprogramming dataset (GSE54848)^7^ where fibroblast cells were OSKM-reprogrammed and followed for 49 days, we observed a rejuvenation (i.e., a biological age decrease) using both inflammatory aging clocks (Fig. 1DE), suggesting that our clocks are sensitive to relevant aging interventions, such as OSKM reprogramming and reinforcing the connection of inflammation downregulation and rejuvenation.

### Evaluation of the ISig inflammatory aging clocks in cancer

Next, we also tested the inflammatory clocks in chronic diseases, starting with cancer, as it has an established connection with both aging and inflammation^8^. As the Blood Inflammatory Clock 2 was more accurate than the Blood Inflammatory Clock 1 in predicting healthy test samples, with lower MAE and higher Pearson R (Fig. 1BC), we decided to employ it solely in the following analyses. So we predicted age by using the Blood Inflammatory Clock 2 to samples from cancer tissues, non-cancer tissues of cancer patients (i.e., adjacent normal tissues), and control tissues from healthy individuals of the EWAS Data Hub 39 cancers dataset (Fig. 2A). The age prediction performance highly dropped (MAE = 17.78, and Pearson’s r = 0.36) compared to the original testing set containing healthy blood samples. The dataset contained samples from cancer tissue (n = 5,675), adjacent normal tissue (n = 864), and control tissue (n = 1,155); however, after age, sex, and tissue matching of the three groups, we were left with 69 samples in each group. We observed that the ISig-based age acceleration of cancer tissues was significantly higher compared to the adjacent normal groups (Fig. 2B). We also observed that the age acceleration of the adjacent normal tissue was higher compared to the healthy control tissues, suggesting accelerated epigenetic age as progression steps from normal to cancer development. Furthermore, we tested four state-of-the-art epigenetic aging clocks, the GrimAge, GrimAge2, Hannum, and Horvath2013 clocks as benchmarks (Supplementary Fig. S1). All benchmark clocks showed significantly higher age acceleration of the cancer tissue compared to the adjacent normal tissue (all except the Horvath2013 clock), and significantly higher age acceleration of the adjacent normal tissue compared to the control tissue (all except the Hannum clock). Thus, ISig is comparable to established clocks in detecting epigenetic age acceleration in the process of carcinogenesis.

**Fig. 2.**
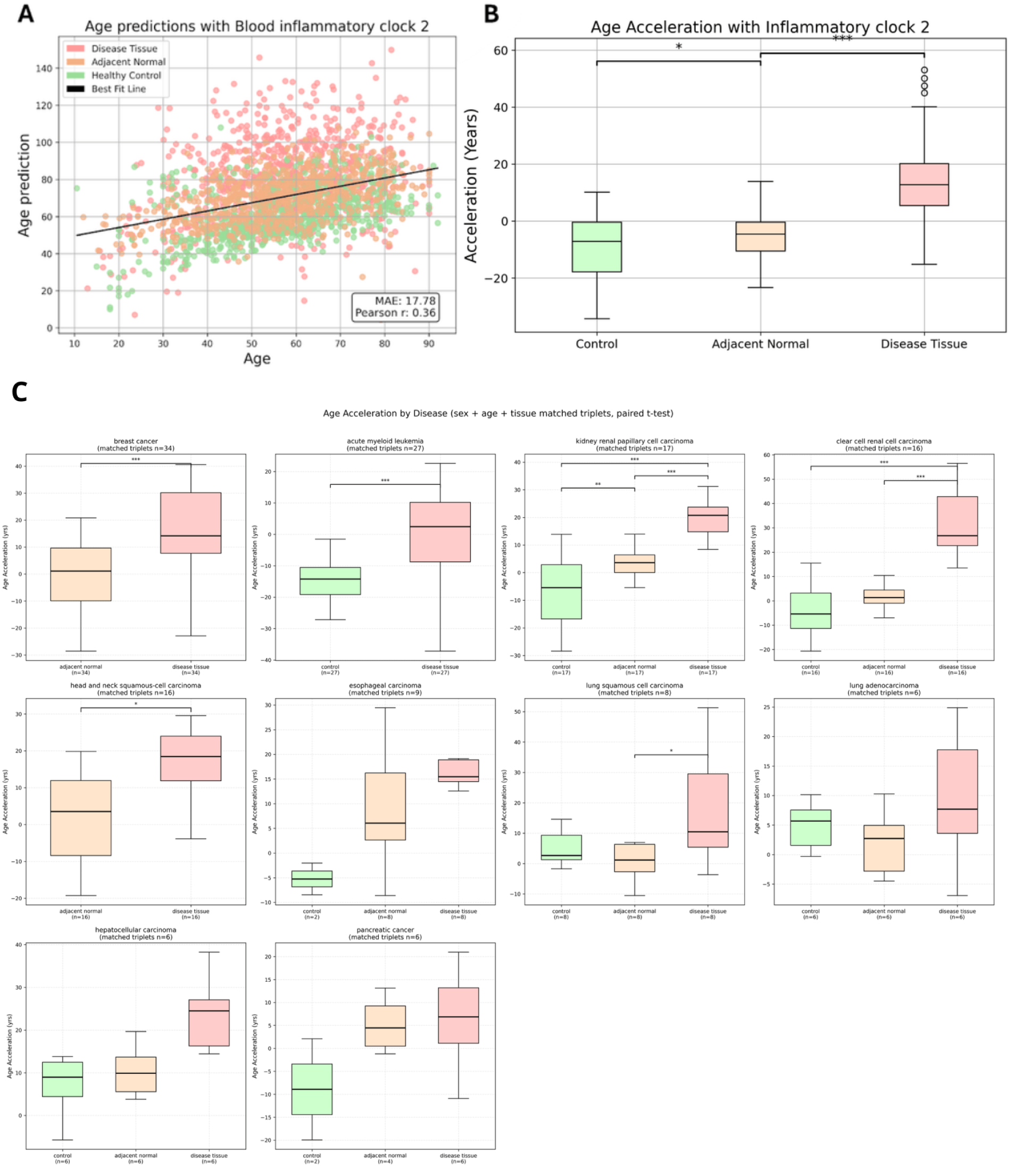
Evaluation of the ISig inflammatory aging clocks in cancer. **A**, Predictions of Inflammatory Clock 2 on the EWAS Data Hub 39 Cancers dataset, and associated accuracy metrics (MAE and Pearson’s r). The black line is the best-fitting line. The dataset contained 5,675 samples from *Disease Tissue*, 864 samples from *Adjacent Normal* tissue, and 1,155 samples from *Healthy Controls*. The three groups are age, sex, and tissue matched; each contains 69 samples (triplets with same sex, tissue and 1.01 years of mean age difference). **B,** Age acceleration (i.e., the difference between the predicted value and the value of the best-fitting line) across three groups: healthy tissues from cancer-free individuals (*Control*), non-cancerous tissues adjacent to tumors (*Adjacent Normal*), and tumor tissues (*Disease Tissue*). After age, sex, and tissue matching, each group contained 69 samples. **C,** The same analysis, but separated by disease.

### Evaluation of the ISig inflammatory aging clock in various non-cancer diseases

The ComputAgeBench benchmarking dataset^9^ includes more than 10,000 blood-derived samples from various sources and cell types such as whole blood (n = 5,790), CD14⁺ monocytes (n = 1,440), and peripheral blood lymphocytes (PBL, n = 1,017). Covering major disease categories (n means the samples left after pair matching) such as immune system disease (ISD, n = 540), neurodegenerative disease (NDD, n = 749), cardiovascular disease (CVD, n = 8), musculoskeletal disease (MSD, n = 466), metabolic bone disease (MBD, n = 20), respiratory system disease (RSD, n = 10), and polygenic score samples (PGS, n = 32). The Blood inflammatory clock 2 was used to predict age using the ComputAgeBench benchmarking dataset (which has no overlap with the ComputAgeBench traning dataset where the clock was trained) (Fig. 3A). We observed significantly increased age acceleration in the case of CVD, ISD, and MSD compared to the healthy controls (Fig. 3B). Agreeing with our ISig-based inflammatory clock, two or three of the four benchmark aging clocks also showed a significant increase for CVD, ISD, and MSD, however, one or two of them showed significant increase of PGS, MDD, and NDD as well (Supplementary Fig. S2).

**Fig. 3.**
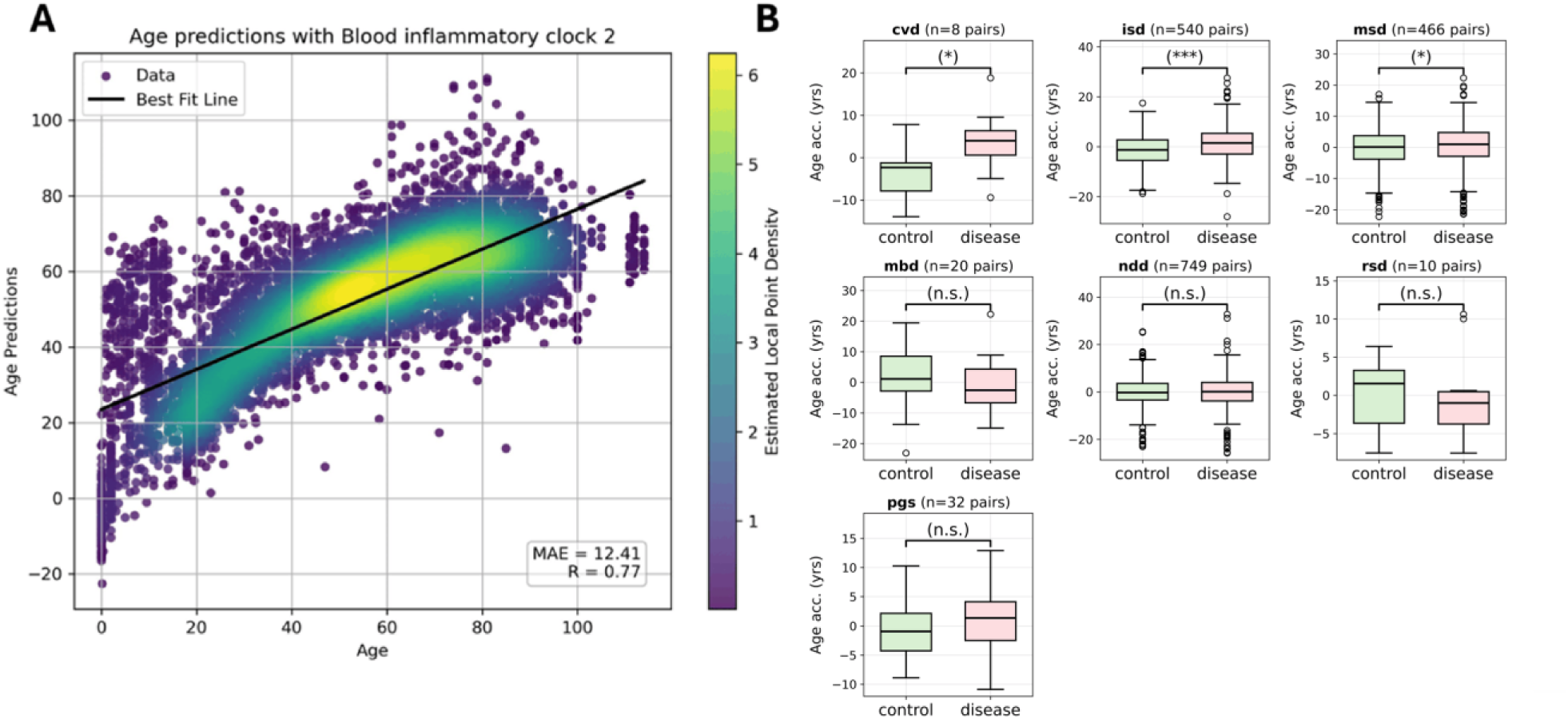
Evaluation of the ISig inflammatory aging clock in various non-cancer diseases. A, Age predictions by using the ISig Inflammatory Clock 2 on the ComputAgeBench benchmarking dataset. **B**, Comparing age accelerations between healthy control and diseased samples of the ComputAgeBench benchmarking dataset. In every compared boxplot, the control and diseased samples originated from the same studies, and blood samples, as well as age, sex, and cell-type matched (pairs matched in the comparison).

### The GeneClock Studio, a web application for the development of gene set-based aging clocks

We developed a user-friendly web application, GeneClock Studio, that enables users to apply the ISig inflammatory aging clocks on external datasets (Fig. 4). Additionally, the platform provides the option of training custom aging clocks tailored to specific targeted gene sets. The CpG sites located on the promoter region of the given genes will be the model features. The GeneClock Studio is available at https://ilab.sztaki.hu/geneclockstudio/).

**Fig. 4.**
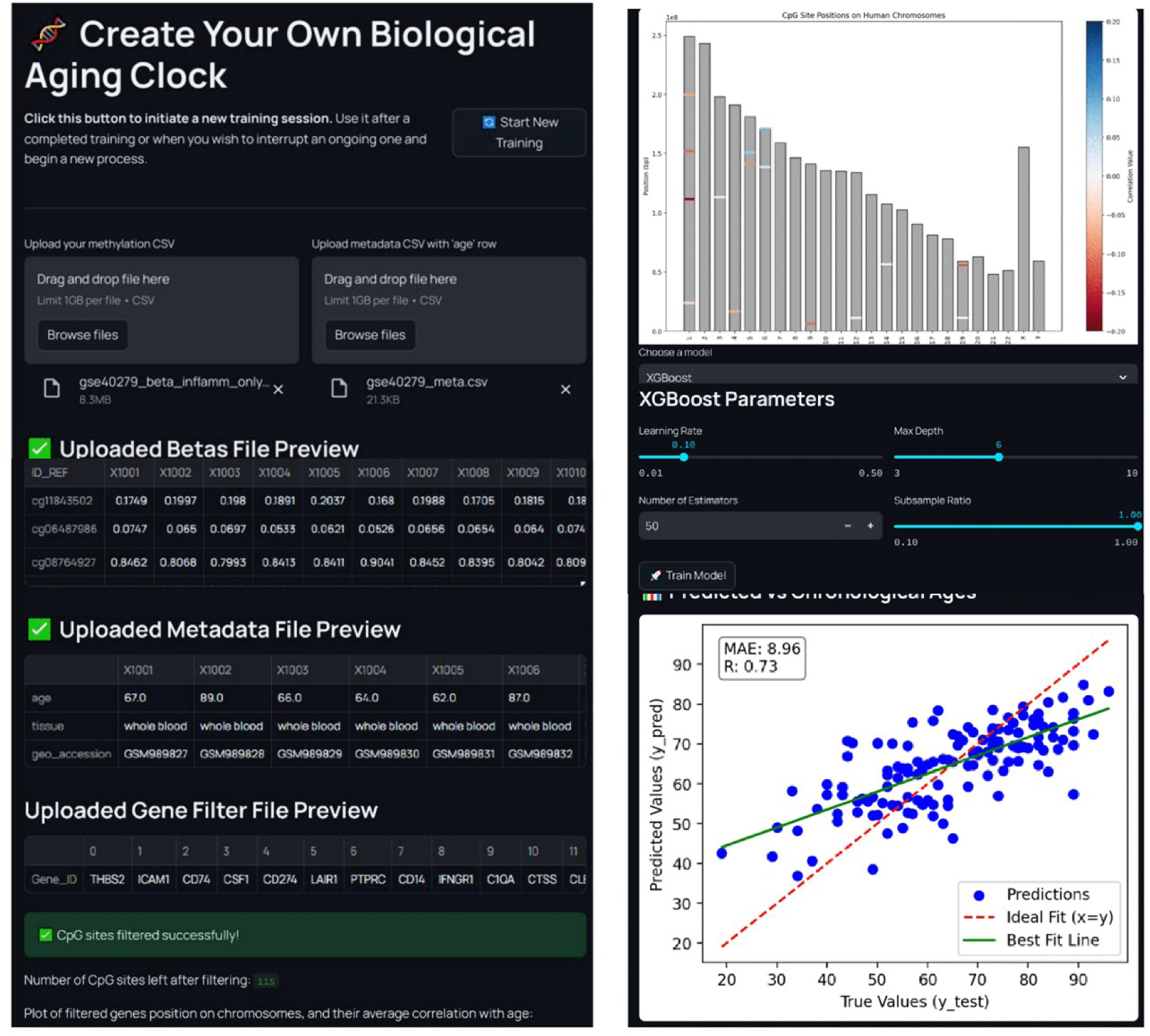
The GeneClock Studio, a web application for the development of gene set-based aging clocks. Screenshots illustrating the training process, beginning with the upload of the meta and beta value data, followed by the selection of a gene filtering method, the upload of a gene filter list, choosing a model, and configuring its hyperparameters. In the web application, users can interactively generate predictions using the inflammatory aging clocks by uploading their data. It is also possible to train a gene set–specific clock without any prior programming knowledge, as the entire process takes place through an interactive interface. Upon gene selection, the application generates a chromosome diagram indicating the location of the selected genes. Color indicates the average correlation of the CpG sites of the gene with age. There are three built-in model options (the ones that usually perform the best on methylation data), ElasticNet, XGBoost, and RandomForest. The hyperparameters of these models are fine-tunable.

## Discussion

This study presents the development and validation of two novel inflammation-based epigenetic aging clocks derived from CpG sites located in the promoter regions of ISig genes. By focusing on a biologically curated inflammatory gene set, these clocks capture a specific and mechanistically interpretable dimension of epigenetic aging, linking inflammatory regulation directly to biological age estimation.

Our findings reinforce the concept that chronic inflammation may represent a central and measurable axis of biological aging encoded in the epigenome as suggested before^10,11^. Unlike conventional epigenetic clocks that rely on large sets of CpG sites selected primarily for predictive performance, our approach demonstrates that a relatively small, functionally coherent subset of the methylome is sufficient to recapitulate aging dynamics. Consequently, our results support the development of pathway-informed epigenetic clocks capable of capturing distinct dimensions of biological aging.

The ISig clocks showed strong predictive performance on healthy samples and were able to detect both accelerated aging in disease and rejuvenation during cellular reprogramming. Notably, in the OSKM reprogramming model, we observed a clear decrease in predicted biological age over time. This finding highlights that the inflammatory epigenetic signatures captured by our models are not static but dynamically malleable. Given that OSKM reprogramming is known to suppress pro-inflammatory gene expression and reset epigenetic marks^12^, our results further support the idea that inflammation is a modifiable driver of epigenetic aging. The sensitivity of the ISig clocks to such interventions suggests their potential utility in evaluating therapies targeting inflammatory pathways and in quantifying rejuvenation effects.

In disease contexts, the ISig clocks revealed expected patterns of epigenetic age modulation. Consistent with the concept of “inflammaging”, we observed increased age acceleration in cancer and immune-related diseases. These findings align with previous reports linking heightened inflammatory activity to accelerated biological aging^10^. In cancer, the observed gradient of age acceleration, highest in tumor tissue, intermediate in adjacent normal tissue, and lowest in healthy controls, further supports the role of inflammation and local tissue environment in shaping epigenetic aging trajectories^15–17^. When compared to established epigenetic clocks such as GrimAge, GrimAge2, Hannum, and Horvath, the ISig-based clocks demonstrate comparable predictive performance while offering increased biological interpretability. In cancer datasets, the ISig clock showed one of the highest differences in age acceleration between adjacent normal tissue and disease samples.

Finally, the development of the GeneClock Studio provides a practical framework for extending this approach. By enabling users to train and apply gene set–specific aging clocks, the platform facilitates exploration of other biological pathways and their relationship to aging. This may accelerate the identification of pathway-specific aging signatures and contribute to a more comprehensive, multidimensional understanding of biological aging.

In summary, our study demonstrates that a curated inflammatory gene set can be used to construct accurate and biologically meaningful epigenetic clocks that capture aging, disease-associated deviations, and rejuvenation dynamics. These findings highlight inflammation as a central, yet distinct, component of biological aging and support the use of pathway-specific clocks to dissect the complexity of aging across different physiological and pathological contexts.

## Methods

### Description and preprocessing of the datasets

The ComputAgeBench dataset (https://huggingface.co/datasets/computage/computage_bench)^9,22^ was collected by combining publicly available DNA methylation array datasets from human blood and saliva samples from the NCBI Gene Expression Omnibus (GEO). Originally, it contained two datasets: (i) the benchmarking dataset comprised 10,410 samples (DNA methylation sites) from 65 studies, while (ii) the training dataset consisted of 7,419 samples from 46 studies (some features are missing in some datasets). The authors selected datasets containing DNA methylation microarrays (27K/450K/850K) samples from blood, saliva, and buccal cells of 18-90 year old subjects covering 19 putative aging accelerating conditions (see Table A1 in ^9^ for the definition of the conditions). In our workflow, we obtained the ComputAgeBench repository using snapshot_download, which stores each sub-study as an individual Parquet file in the ./data/train/ and ./data/benchmark/ directories, accompanied by two metadata files (computage_train_meta.tsv and computage_bench_meta.tsv). Each Parquet file was loaded separately, column names were harmonized to lowercase, and missing beta values were imputed using the per-CpG column mean. Beta values were additionally validated to ensure they fell within the expected [0, 1] range. After discarding low-quality samples (more than 50% missing value), the ComputAgeBench training dataset contained 6,039 samples and the benchmark dataset 9,841 samples with an approximately normal age distribution ranging from 0 to 99 years, and most samples concentrated between the ages of 30 and 60 years.

The EWAS Data Hub (https://ngdc.cncb.ac.cn/ewas/datahub)^23^ is a public repository that integrates DNA methylation array data (including 450K, 850K, and 935K in the current version) and comprehensive metadata from various data sources (GEO, TCGA, ENCODE, and ArrayExpress) based on a standard pipeline to minimize batch effects from distinct sources. The EWAS Data Hub provides DNA methylation profiles of 136 diseases. For each disease, the methylation profiles of disease samples, as well as gender and age-matched controls, are provided. From the download page of the EWAS Data Hub ((https://ngdc.cncb.ac.cn/ewas/datahub/download), we acquired the “DNA methylation profiles of 39 cancers” dataset (referred to as *EWAS Data Hub 39 cancers* datasets). We discarded the samples of the dataset GSE64491 with 30 different tissue samples from a 112-year-old woman.

### Development of the ISig-based inflammatory aging clocks

We trained aging clocks on the CpG sites of the promoter regions (TSS1500, TSS200, 5’UTR, and 1stExon) of the ISig genes since they had an instrumental role in the regulation of DNA replication and transcription. The inflammatory aging clocks were trained using an ElasticNet regression model implemented via a MinMax-scaled ElasticNet pipeline, with hyperparameters optimized through a 5-fold cross-validated grid search. The parameter grid spanned ElasticNet α values ranging from 0.0001 to 1 (20 linearly spaced values) and l1-ratio values from 0.1 to 0.9 (10 linearly spaced values). The models were trained exclusively on healthy samples. The Inflammatory clock 1 was trained on the Hannum et al. (GSE40279) dataset^24^, and the Inflammatory Clock 2 was trained on the cleaned ComputAgeBench training set^9^ with 6039 samples. An 80:20 train–test split was applied to partition the datasets. The clock performance was measured by mean absolute error (MAE) and Pearson’s correlation coefficient (R). For Inflammatory clock 2, training with ElasticNet was also superior to XGBoost: while results on the 20% held-out portion of the ComputAgeBench training set were similar (ElasticNet: MAE = 5.9 years, R = 0.94; XGBoost: MAE = 5.4 years, R = 0.93).

### Application of the ISig-based inflammatory aging clocks

We filtered the application datasets based on the ISig gene set. The TSS1500, TSS200, 5’ untranslated region (5’UTR), and the first exon (1stExon) regions were considered promoter regions. During preprocessing, missing values are imputed with the average methylation level of the CpG sites. However, any samples or features with too many missing values (more than 50%) are removed to maintain reliable results. We did not separate studies of the ComputAgeBench, which can correspond to multiple studies.

### Application of the benchmark aging clocks

We applied available state-of-the-art epigenetic aging clocks as benchmarks, such as the Hannum^24^, Horvath2013^25^, GrimAge^26^, and GrimAge2^27^ clocks. We used the pyaging Python library^28^. For each clock, DNA methylation beta values were transposed into a sample-by-CpG matrix, converted to AnnData format, and age predictions were generated using pya.pred.predict_age. Any CpG sites required by a clock but absent from the dataset were imputed using the mean strategy built into pyaging’s df_to_adata function; for GrimAge specifically, the sex and age were also added.

### Age acceleration

Age acceleration was calculated as the difference between the predicted values and the values of the best-fitting line, which was fitted using the samples included in each respective comparison.

### Development of the GeneClock Studio

The web application is built on a Python-based client–server architecture with a FastAPI REST backend and a Streamlit multi-page frontend. The backend is served via Uvicorn over HTTPS (self-signed SSL) and is organized into four routers: authentication, dataset management, aging clock prediction, and model training, each backed by dedicated service modules. User access is controlled via JWT-based authentication (Passlib/python-jose). The frontend communicates with the backend through HTTP requests carrying Bearer tokens, and exposes three functional pages: dataset inspection, application of built-in epigenetic aging clocks, and custom clock training. The training pipeline supports ElasticNet, XGBoost, and Random Forest models (scikit-learn/XGBoost) fitted on user-supplied DNA methylation data, optionally restricted to an uploaded gene set, with trained models persisted server-side in a JSON-tracked registry and made available for download.

### Statistics

Statistical comparisons between groups were performed using Welch’s two-sample t-test (scipy.stats.ttest_ind, equal_var=False) to account for unequal variances. Significance thresholds were set at p < 0.05 (*), p < 0.01 (**), and p < 0.001 (***); non-significant results are labelled n.s. Clock performance was evaluated using Mean Absolute Error (MAE = mean of |predicted − chronological age|) and Pearson correlation coefficient (R), both computed with NumPy and SciPy.

### Data and Code Availability

We used the publicly available Hannum et al. dataset (GSE40279), Ohnuki et al. dataset (GSE54848), ComputAgeBench dataset (https://huggingface.co/datasets/computage/computage_bench)^9,22^, and the EWAS Data Hub “DNA methylation profiles of 39 cancers” dataset (https://ngdc.cncb.ac.cn/ewas/datahub/download). For the gene signature used, please contact the corresponding authors, who may provide the signature upon reasonable request.

### Sex, age, and tissue matching

In every boxplot comparison, the samples were age, sex, and tissue matched (some already contained the same tissue samples). We created pairs (or triplets in the caner comparison) with the same sex, tissue and minimal age difference but maximum 2 years across the pairs.

## Acknowledgements

The project was supported by the HUN-REN (TKCS-2024/37), and European Union project RRF-2.3.1-21-2022-00004 within the framework of the Artificial Intelligence National Laboratory and supported by the DecodAge project (COMPETE2030-FEDER-00651300, Project nr. 15507) funded by Portugal 2030 and co-funded by the European Union. CK was also supported by the National Research, Development and Innovation Office – NKFIH (FK-146113), and the János Bolyai Research Scholarship of the Hungarian Academy of Sciences.

## Supplementary Figures and Tables

**Supplementary Fig. S1.**
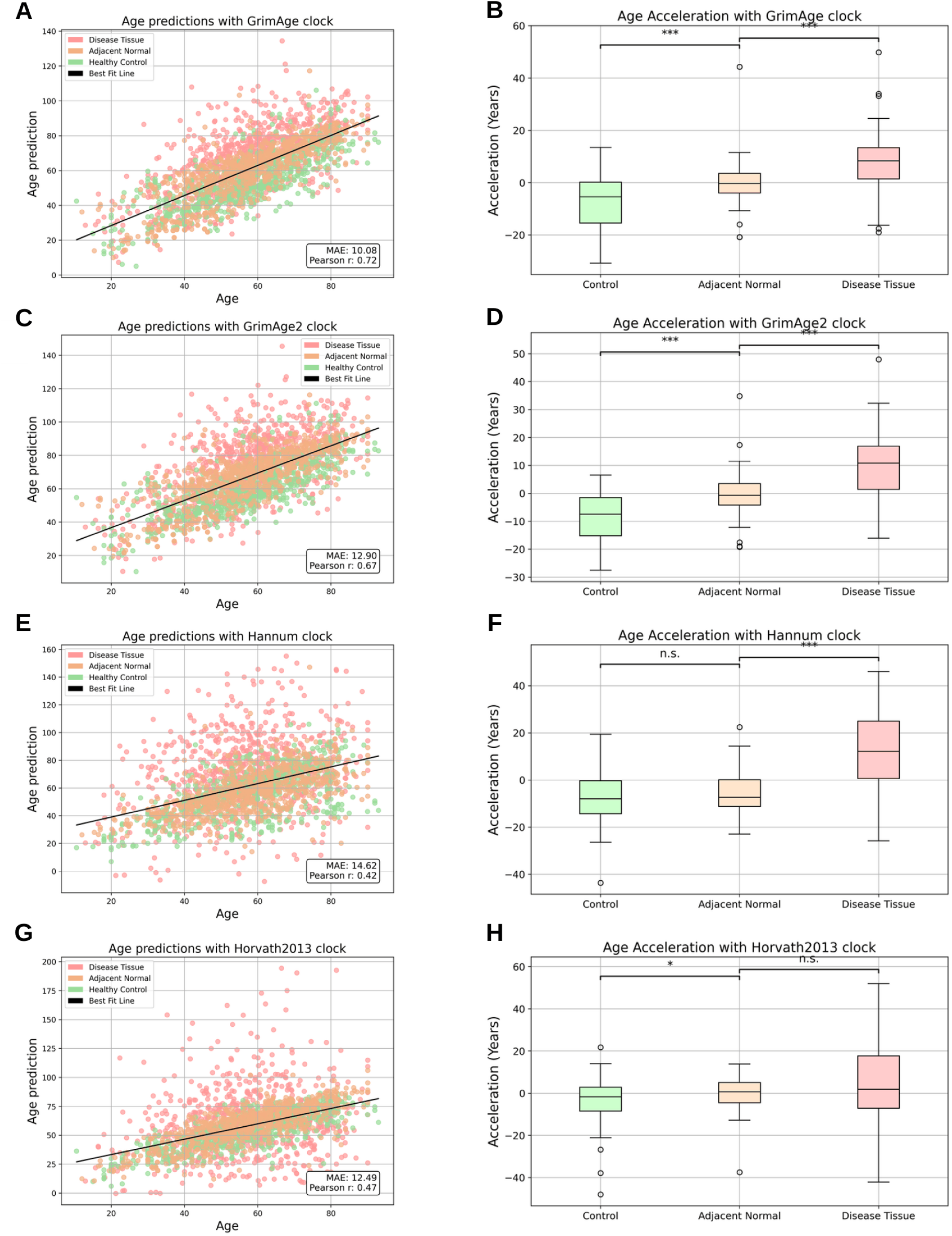
Evaluation of the benchmark epigenetic aging clocks in cancer. The same analysis as in Fig. 3, but here we applied the **AB,** *GrimAge clock*, **CD,** *GrimAge2 clock,* and **EF**, *Hannum clock, **GH,** Horvath2013 clock,* using the EWAS Data Hub 39 Cancers dataset. In each boxplot there are 2271 samples. In all comparisons, control and diseased, and were consistently matched for age, sex, and tissue (we found 39 of triplets with the same sex, age, and tissue across every comparison and in all comparisons we were left with 69 samples in each group).

**Supplementary Fig. S2.**
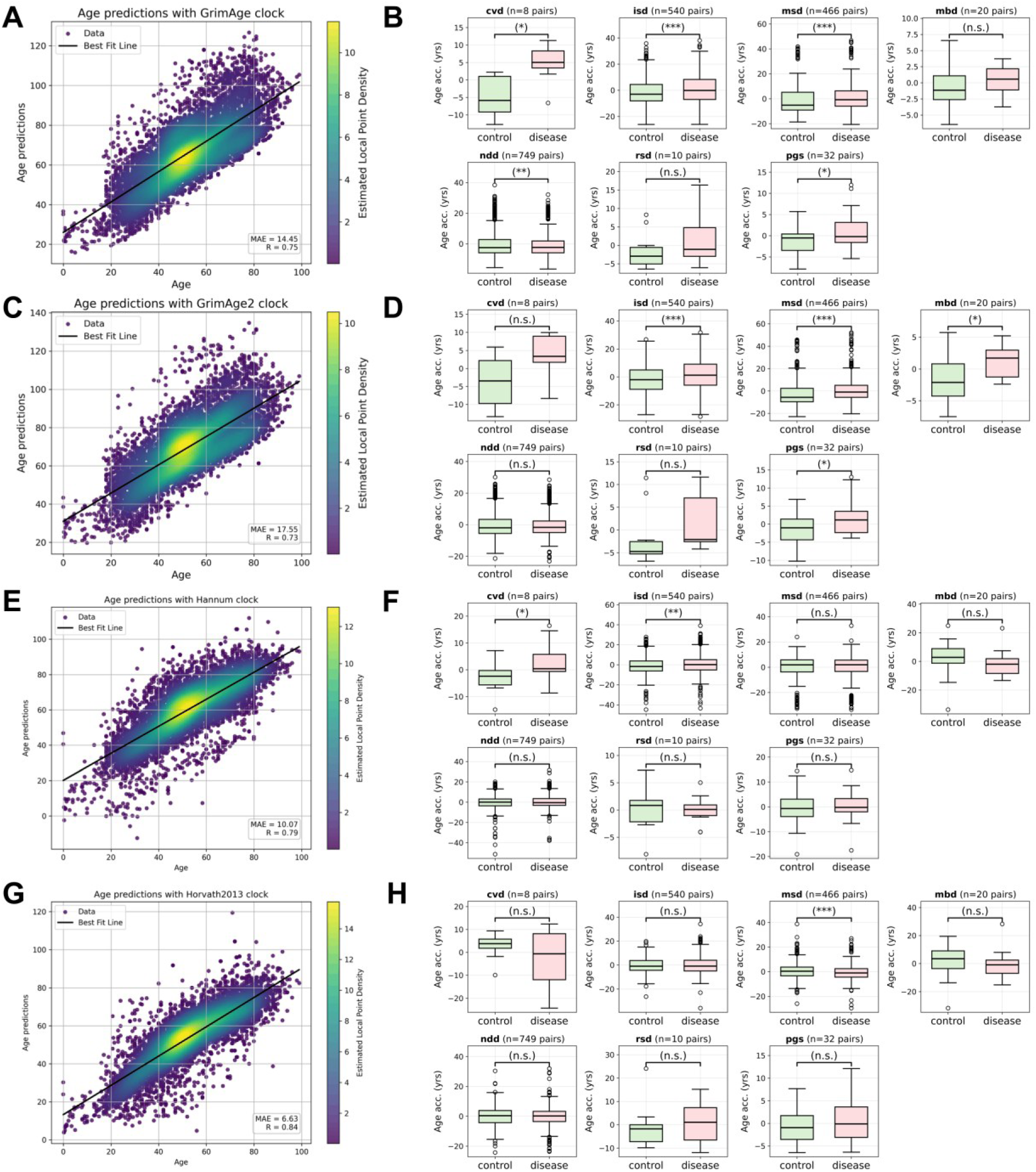
Evaluation of the benchmark aging clocks in various diseases of the ComputAgeBench dataset. The same analysis as in Fig. 4, but here we applied the **AB,** *GrimAge clock* **CD,** *GrimAge2 clock*, and **EF**, *Hannum clock,* **GH** *Horvath2013 clock*, on the non-cancer blood samples of the ComputAgeBench dataset. In all comparisons, control and diseased samples came from the same studies, and were consistently matched for age, sex and cell-type (we created pairs with same sex, age and cell type across every comparison).

